# DeconV: Probabilistic Cell Type Deconvolution from Bulk RNA-sequencing Data

**DOI:** 10.1101/2023.12.07.570524

**Authors:** Artur Gynter, Dimitri Meistermann, Harri Lähdesmäki, Helena Kilpinen

## Abstract

Bulk RNA-Seq remains a widely adopted technique to profile gene expression, primarily due to the persistent challenges associated with achieving single-cell resolution. However, a key challenge is accurately estimating the proportions of different cell types within these bulk samples. To address this issue, we introduce DeconV, a probabilistic framework for cell-type deconvolution that uses scRNA-Seq data as a reference. This approach aims to mitigate some of the limitations in existing methods by incorporating statistical frameworks developed for scRNA-Seq, thereby simplifying issues related to reference preprocessing such as normalization and marker gene selection. We benchmarked DeconV against established methods, including MuSiC, CIBERSORTx, and Scaden. Our results show that DeconV performs comparably in terms of accuracy to the best-performing method, Scaden, but provides additional interpretability by offering confidence intervals for its predictions. Furthermore, the modular design of DeconV allows for the investigation of discrepancies between bulk-sequenced samples and artificially generated pseudo-bulk samples.

## 1. Introduction

Bulk RNA-sequencing (bulk RNA-seq) is the main source of transcriptomic data (Supplementary methods), as it is a mature technology that is generally more available and cost-efficient than single-cell RNA-sequencing (scRNA-seq). However, changes of gene expression observed between heterogeneous bulk RNA-seq samples can be driven by two distinct phenomena: differential regulation of gene expression within cells or changes in cell types relative proportions. This can lead to occurrences of Simpson’s paradox (Trapnell 2015) where false conclusions can be deducted from differential expression analysis. Usage of scRNA-seq has also shown that some cell types can not be identified with a small set of markers and therefore need a high-troughput profiling (Ianevski et al. 2022). However, scRNA-seq is tied by its cost, especially in experimental designs that requires comparing multiple population of individuals (i.e. reference vs target condition studies). In that case, the number of profiled cells required to represent cell type diversity and proportion of each individual increases dramatically. Another dignificant challenge is that tissues cannot always be easily dissected by scRNA-seq, as some contains cell types that are hard to dissociate and increase the time, cost and effort necessary to prepare the scRNA-seq library. Hence, bulk RNA-seq in combination with a deconvolution method to infer cell type proportions offers an attractive alternative to scRNA-seq.

Reference-based deconvolution techniques leverage a predefined gene expression dataset to infer the proportional abundance of distinct cell types within a target bulk RNA-seq sample. Such a reference represents the cell types anticipated to be present in the target sample. It may originate either from bulk RNA-seq where each sample corresponds to a cell type, or more usually scRNA-seq. Notably, the use of single-cell RNA-seq data as the reference confers a higher degree of precision, thereby enhancing the overall accuracy of the deconvolution process. Comparative evaluations have demonstrated that reference-based deconvolution methods generally outperform both marker gene-based and other non-reference-based strategies, as evidenced by a recent benchmark study (Jin and Liu 2021). Consequently, when gene expression profiles of the anticipated cell types are accessible, reference-based deconvolution serves as the method of choice.

While existing state-of-the-art deconvolution techniques, such as CIBERSORTx (Newman et al. 2019), MuSiC (Wang et al. 2019), and Scaden (Menden et al. 2020) have advanced the field and provided valuable insights, they also present certain limitations as as they do not permit to quantify the uncertainty of their predictions. Moreover, they do not use the advancement in probabilistic modelling of RNA-Seq count (Risso et al. 2018) and hence present a complex methodology to achieve their task. In contrast, our proposed framework, DeconV, adopts a simple, probabilistic and modular approach that leverages advanced statistical paradigms developed for RNA-Seq analysis, thereby addressing these limitation while maintaining high levels of accuracy and speed.

## 2. Results

### 2.1. Method overview

Similarly to MuSiC (Wang et al. 2019), DeconV assumes a linear-sum-property between single-cell and bulk gene expression, implying that bulk gene expression is a sum of the components from single-cell gene expression. In contrast to MuSiC, DeconV models cell-type-specific gene expression with probability distributions (**Figure 1**) as opposed to point estimates. This is motivated by several biological and technical factors; First, cells of a same type are not identical. They can vary in size and their processes are regulated based on signals coming from the environment as well as other cells. Thus, gene expression (and messenger RNA (mRNA) quantity in cells) often fluctuates depending on the state of the cell. For this reason, single-cell gene counts are often over-dispersed, i.e. the standard deviation is often larger than the mean (Love et al. 2014). Secondly, scRNA-seq (and bulk RNA-seq) measurements are not perfect due to technical limitations. For instance, dropout is a unique phenomenon in single-cell gene expression where certain mRNA molecules of active genes are not detected which results in loss of information and can lead to biased estimation of cell-type-specific gene expression. Such technological limitations introduce noise to gene expression measurements and therefore we hypothesise that using probability distributions instead of point estimates helps to account for these factors and improve the accuracy of cell type deconvolution. We also choose to make DeconV modular, as the user can choose among five distributions (**Table S1**) to model gene expression and whether or not to account for zero-inflated distribution (**Figure 1c**).

**Figure 1:**
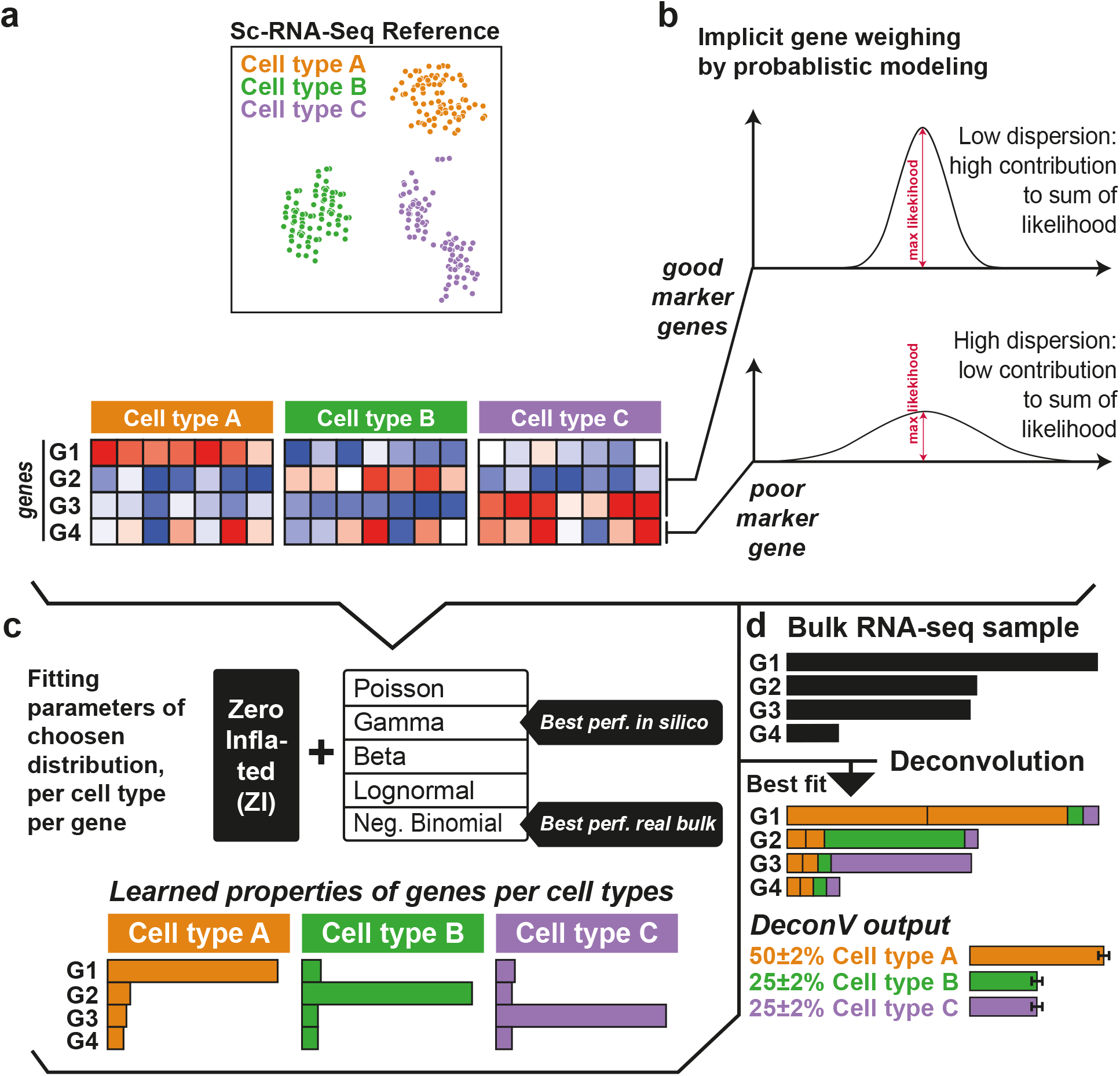
Overview of DeconV methodology. **a**. scRNA-seq example of a reference dataset, containing three cell types. Three marker and one non informative genes are represented in the heatmap. **b**. The probabilistic framework of DeconV leads to an implicit weighing of genes during deconvolution, as the low dispersion of marker genes in their cell type will have a stronger effect during the fitting phase than a non informative gene. **c**. Parameters of the chosen distribution are then learned from the reference dataset. Benchmarks show that zero-inflated Gamma distribution is performing best with *in silico* bulk RNA-seq reconstructed from scRNA-seq but zero-inflated negative binomial has better results with real bulk RNA-Seq. **d**. Cell-types proportions are estimated from the query dataset after maximizing the likelihood of genes and computing the best fit. In this schematic, there is two times more cell type A than B or C. DeconV also provides the confidence intervals (default at 95%) of the estimated proportion, in this example 2% for each cell type.

Using probability distributions to model gene expression enables three further improvements to the deconvolution; First, a loss function with a probabilistic interpretation can be used as opposed to a commonly used loss function based on squared error, e.g. non-negative least squares (NNLS) used by MuSiC. This allows to measure the distance between pseudo-bulk and real bulk gene expression (**Figure 1b**) in proportional manner, equally across the whole range of lowly- and highly-expressed genes. On the other hand, the sum of NNLS is dominated by the distance between pseudo-bulk and real bulk of highly expressed genes. Secondly, using probability distribution allows to implicitly up- and down-weigh the effect of certain genes on deconvolution. For example, a noisy gene (e.g. gene ‘*G4*’ in **Figure 1a**) provides little to no information to distinguish between cell types while good marker genes (e.g. genes *‘G1’, ‘G2’, ‘G3’*) are essential for cell type deconvolution. Therefore, up-weighing consistent marker genes and down-weighing non-informative highly dispersed genes can improve the accuracy of cell type deconvolution. Probability density (or mass, in case of discrete random variable) of highly dispersed gene is spread across a larger range than of a consistent marker gene. Therefore, a higher maximum likelihood can be achieved in good marker genes (**Figure 1b**) which ultimately up-weights them when optimizing the likelihood function. In contrast to DeconV, MuSiC implements gene weighing with a heuristic function based on single-cell gene-specific variance (Wang et al. 2019). Finally, the cell type proportions are estimated as probability distributions as opposed to point estimates like in most if not all state-of-the-art methods. This allows to quantify the uncertainty of a prediction, e.g. by specifying a 95% confidence interval.

On the other hand, deep-learning-based methods, e.g. Scaden (Menden et al. 2020), have shown a great potential to accurately predict cell type proportions from bulk RNA-seq. However, due to the scarcity of real bulk datasets with known cell type proportions, these methods are generally trained exclusively on artificially pseudo-bulk datasets generated from a single-cell reference. Thus, the model has never encountered real bulk gene expression data prior to deconvolution. This raises concerns whether the results of these models can be trusted. Moreover, so called black-box design of the model with no estimation of uncertainty makes the interpretation of the results difficult.

### 2.2. Benchmark

DeconV was benchmarked with two independent dataset (Xin et al. 2016; Monaco et al. 2019, see Material and Methods - Description of benchmarking datasets for more details), along with MuSiC, Scaden and CIBERSORTx. The results are visualised in **Figure 2**. DeconV achieved the lowest estimation error across the cell types in both datasets. In the Xin-dataset (Xin et al. 2016, **Figure 2a**), DeconV (gamma), MuSiC, and Scaden achieved good results indicated by low estimation error (*<* 10%) with DeconV exhibiting the lowest error and the highest correlation by a slight margin (**Table 1**). On the other hand, in the PBMC-dataset, the deconvolution accuracy of all methods decreased (higher error and lower correlation) by a significant margin and the difference between the methods was more pronounced. This was usually due to an over-estimation of a single cell type which led to underestimation of other cell types. For instance, Scaden (orange) overestimated CD4 T lymphocytes (CD4 T), CIBERSORTx (red) overestimated Dendritic Cells (DCs), and MuSiC overestimated Natural Killer Cells (NK cells) (**Figure 2b**). Furthermore, all methods overestimated proportions of rare DCs, highlighting challenges of rare cell type deconvolution, perhaps due to a low signal-to-noise ratio.

**Table 1:** Benchmark summary of DeconV (Gamma or Negative Binomial model) for two independent datasets and with three deconvolution methods from literature.

| Method | Dataset | Correlation (R) | Root Mean Square Error (RMSE) | Mean Absolute Error |
| --- | --- | --- | --- | --- |
| DeconV (Gamma) | Xin-dataset | **0.97** | **0.03** | **0.04** |
| DeconV (NB) | Xin-dataset | 0.64 | 0.20 | 0.15 |
| MuSiC | Xin-dataset | 0.94 | 0.10 | 0.06 |
| Scaden | Xin-dataset | 0.96 | 0.08 | 0.06 |
| CIBERSORTx | Xin-dataset | 0.78 | 0.22 | 0.16 |
| DeconV (Gamma) | PBMC-dataset | 0.05 | 0.15 | 0.12 |
| DeconV (NB) | PBMC-dataset | 0.56 | **0.11** | **0.10** |
| MuSiC | PBMC-dataset | -0.12 | 0.33 | 0.23 |
| Scaden | PBMC-dataset | **0.63** | 0.17 | 0.14 |
| CIBERSORTx | PBMC-dataset | -0.54 | 0.24 | 0.19 |

**Figure 2:**
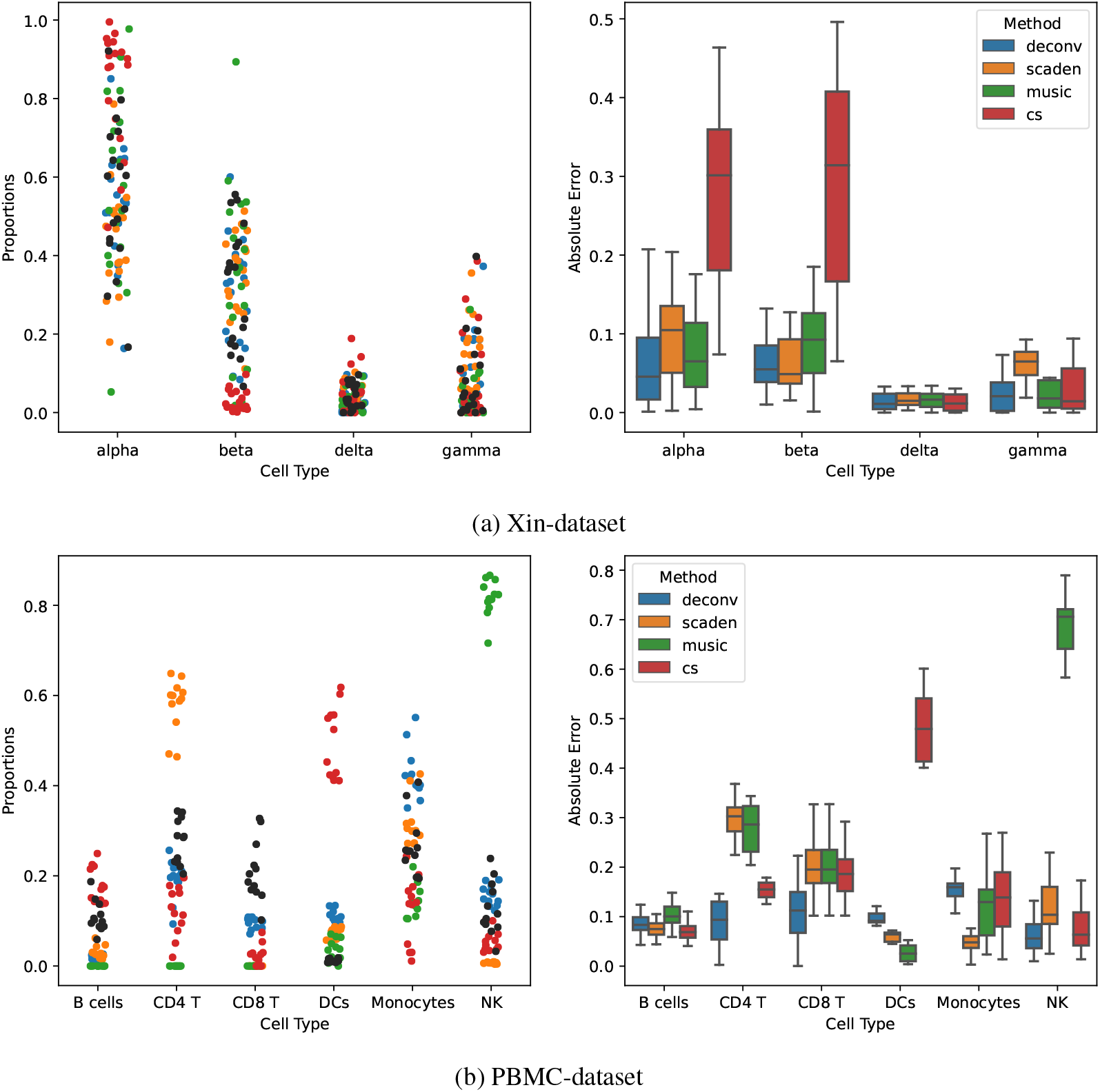
Deconvolution results summary per cell type and method. **a**. Results for the Xin-dataset, where the query dataset is pseudo-bulk generated from the single-cell reference. **b**. Results for the PBMC-dataset. **Left**: Each point represents one-sample cell type proportions. Color represents a deconvolution method. Black points are cell type proportions ground truth validated with flow cytometry. **Right**: Box plot of absolute estimation error per cell type and method. DeconV performs consistently across both datasets.

Deconvolution of the PBMC-dataset (Monaco et al. 2019) was significantly more difficult than the Xin-dataset, as indicated by larger estimation errors (right column of **Figure 2**). While the PBMC-dataset is inherently more complex, due to more cell types as well as smaller and more balanced cell type proportions in general, we hypothesized that the drop-off in accuracy between the two datasets is rather explained by the intrinsic difference between real and pseudo-bulk RNA-Seq. Thus, in the Xin-dataset, the query bulk RNA-seq dataset is fact pseudo-bulk computed from the scRNA-seq, whereas the PBMC-dataset contains real bulk RNA-seq gene expression. These technologies inherently differ in how mRNA-molecules are sequenced which ultimately affects gene expression counts. For instance, paired-end sequencing is commonly used for bulk RNA-seq while scRNA-seq typically utilizes single-end sequencing. This was confirmed by generating a pseudo-bulk from the PBMC-dataset and deconvoluting it with DeconV: the estimated proportions were linearly correlated with the ground truth proportions of the artificial pseudo bulk samples (**Figure S1**).

**Figure 3** shows the results of deconvolution estimations with respect to the ground truth cell type proportions from two datasets and two models, gamma- and negative-binomial-model. Again, the results between the two datasets are inconsistent: gamma-model achieved better results than negative-binomial-model in the artificial pseudo-bulk dataset and vice versa in the PBMC-dataset with real bulk. Notably, negative-binomial-model (**Figure 3b**) exhibited a rather poor results compared to gamma-model (**Figure 3a**) in the Xin-dataset. However, a linear relationship (R=0.64), although with a larger error (RMSE=0.20), is observable from the estimated proportions of negative-binomial-model and ground truth proportions (**Figure 3b**), perhaps suggesting an overestimation of variance of the artificial pseudo-bulk data.

**Figure 3:**
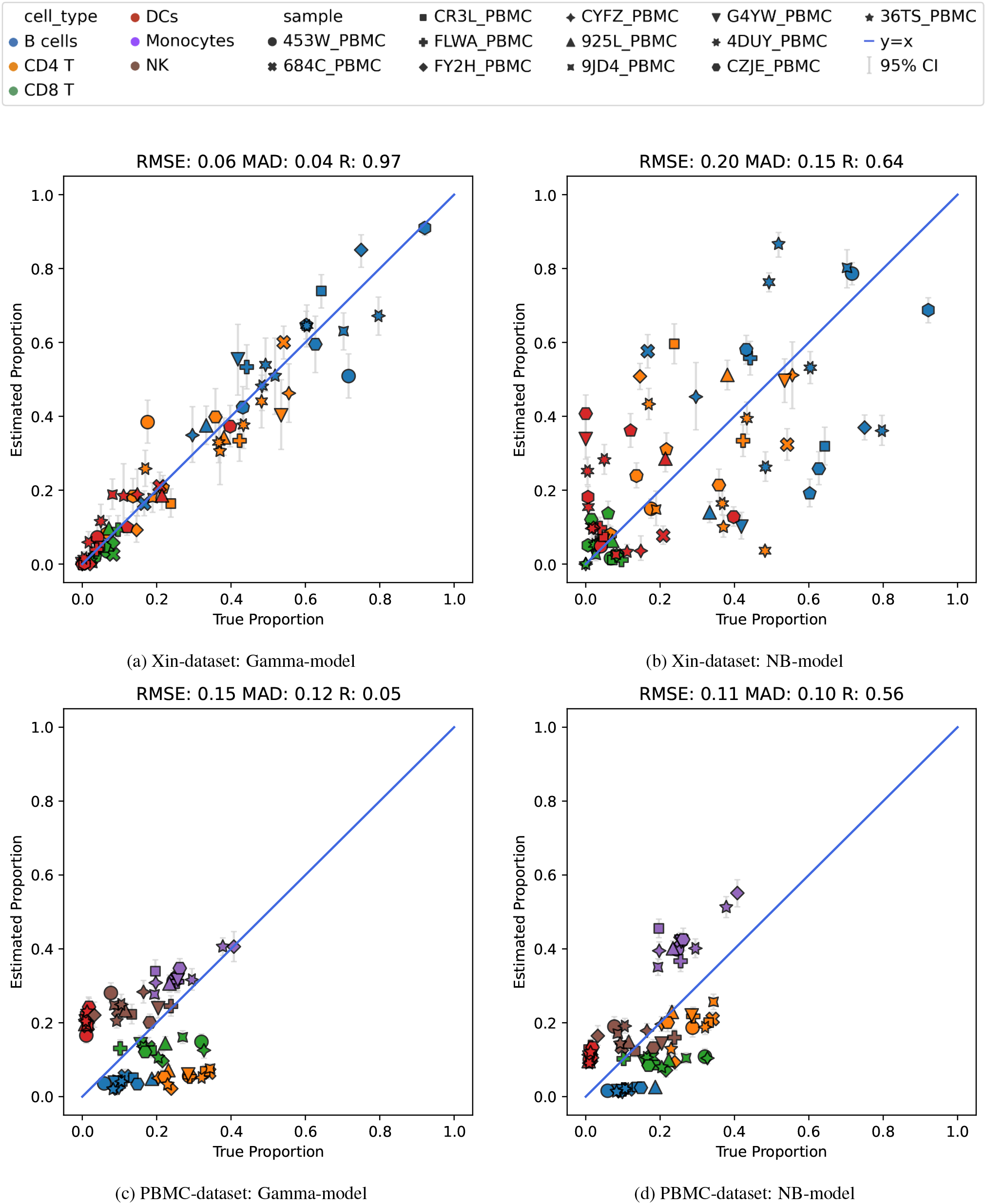
PBMC-dataset estimated (y-axis) vs. ground truth (x-axis) cell type proportions from gamma models (**a**,**c**) and NB-models (**b**,**d**) in pseudo-bulk Xin (**a**,**b**) and in PMBC real bulk RNA-seq datasets (**c**,**d**). Each point corresponds to a cell type proportion in a sample. Shape of the point indicates the sample while color indicates the cell type. 95% confidence interval is highlighted with grey error bars for each point.

## 3. Material and Methods

DeconV consists of two models, a reference model (**Figure 1c**) and a deconvolution model (**Figure 1d**). Reference model learns latent parameters from single-cell reference after which deconvolution model uses the learned parameters to infer optimal cell type composition of a bulk sample.

### 3.1. Reference Model

First, the reference model is fitted using single-cell reference by maximizing the likelihood of the observed data (single-cell gene counts) with respect to the parameters of the reference model (**Figure S2**). The reference model, is a probabilistic model consisting of a discrete distribution (zero-inflated Poisson or zero inflated negative-binomial) with cell-type-specific parameters for single-cell gene counts:

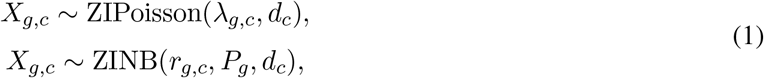

where *X*_*g,c*_ denotes the expression of gene *g* in cell type *c* that is observed from the single-cell reference. Variables *λ*_*g,c*_, *r*_*g,c*_ and *P*_*g*_ are latent random variables assumed to have prior distributions: gamma, log-normal, and beta, respectively (**equation 2; Figure S2**). Parameter *d*_*c*_ denotes the cell-type specific dropout. Since the Poisson distribution underestimates the variance for over-dispersed random variable, such as single-cell gene expression counts, including a compound distribution (gamma-Poisson compound distribution) allows to model over-dispersed random variables. In fact, gamma-Poisson has been previously used to model gene expression (Ahlmann-Eltze and Huber 2020; Thygesen and Zwinderman 2006) and corresponds to the negative-binomial distribution (Greenwood and Yule 1920). While it is possible to model over-dispersed counts directly with the negative-binomial distribution, we assume prior distributions to account for uncertainty and to allow quantification of posterior distribution:

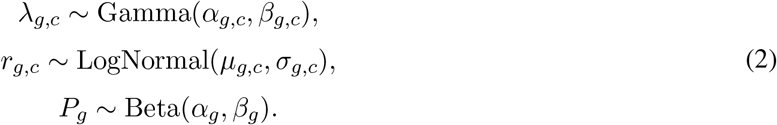

Hyperparameters *α*_*g,c*_, *β*_*g,c*_, *μ*_*g,c*_, *σ*_*g,c*_, *α*_*g*_, *β*_*g*_ and *d*_*c*_ are learned such that the likelihood of the observed data is maximized. This process can be divided into four steps. First, a batch is sub-sampled from the reference single-cell dataset. Second, latent variables *λ*_*g,c*_, *r*_*g,c*_ and *P*_*g*_ from prior distributions are drawn. Third, the log-likelihood of the model conditional on the sampled values of the latent variables is calculated with respect to the observed data, i.e. batch of single-cell reference. Finally, backpropagation, i.e. optimization of the prior hyperparameters *α*_*g,c*_, *β*_*g,c*_, *μ*_*g,c*_, *σ*_*g,c*_, *α*_*g*_, *β*_*g*_ and *d*_*c*_ is carried out by taking the gradient of log-likelihood. This process is repeated until the convergence or specified maximum number of iterations is reached. This results in optimized hyperparameters 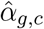,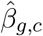,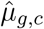,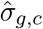,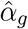,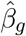, and 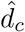.

### 3.2. Deconvolution Model

Once the reference model has been fitted, deconvolution model translates single-cell expression to pseudo-bulk or real bulk gene expression. For pseudo-bulk data, this is motivated by the aggregation-property of Poisson distributions which states that the sum of two (or more) Poisson random variables has also a Poisson distribution. For pseudo-bulk or real bulk gene expression count *B*_*g*_ for gene *g*, the model using the Poisson distribution is defined as:

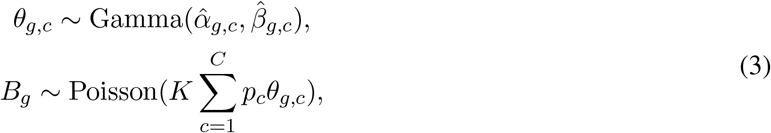

where variable *K* is a scaling factor which is proportional to the number of cells sequenced and the sequencing depth of the bulk sample, and variable *p*_*c*_ is the proportion of cell type *c*. Cell type proportions (*p*_1_, …, *p*_*C*_) are modeled with the Dirichlet distribution:

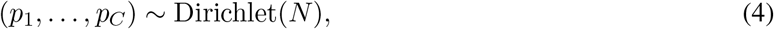

where *N* is a hyperparameter with the size equal to the number of cell types and is initialized as ones (uninformative prior). The model using the negative binomial is defined similarly:

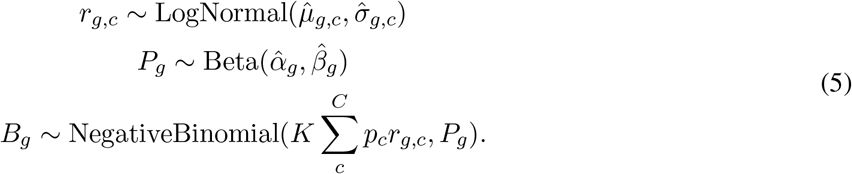

The hyperparameter *N* is optimized similarly as the reference model. First, proportions are drawn from prior distribution (Dirichlet **eq. 4**). Second, the log-likelihood of the observed bulk RNA-seq count data is calculated conditional on the sampled latent variables and by using the optimized hyperparameters 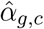, 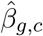,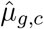,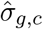,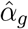, and 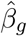. Finally, the optimization of hyperparameter *N*, which defines the distribution for bulk cell type proportions, is carried out by backpropagation of the log-likelihood gradient.

In contrast to scRNA-seq, any possible dropout in bulk RNA-seq is convoluted behind the sum of gene expressions. Thus, dropout is not an observable value from bulk gene expression counts. The extent of dropout caused by technical factors in sequencing compared to a biological dropout is unknown. Biological dropouts are expected to be cell type specific and similar between scRNA-seq and bulk RNA-seq, whereas technical dropouts originate from differences in the underlying sequencing technologies. DeconV implements five ways to modeling dropout, listed in **Table S1**. Dropout is implemented as a deterministic parameter and is optimized during single-cell-reference fitting.

Dropout is a technical phenomenon common to scRNA-seq. In order to model dropout, DeconV uses zero-inflated distributions for gene expression counts. This adds an additional parameter to the model which denotes the probability of observing zero. DeconV implements two types of dropouts: one which is shared across cell types *d*_*g*_ and one which is unique for each cell type *d*_*g,c*_. In case of a separate dropout, gene expression dropout in bulk, which if exists is hidden behind the sum of gene expression, is calculated using proportionate dropout:

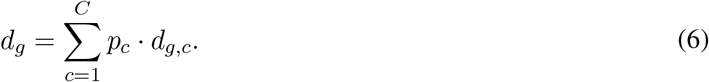

In contrast to scRNA-seq, any possible dropout in bulk RNA-seq is convoluted behind the sum of gene expressions. Thus, dropout is not an observable value from bulk gene expression counts. The extent of dropout caused by technical factors in sequencing compared to a biological dropout is unknown. Biological dropouts are expected to be cell type specific and similar between scRNA-seq and bulk RNA-seq. whereas technical dropouts originate from differences in the underlying sequencing technologies. Dropout estimation is implemented as a deterministic parameter and is optimized during single-cell-reference fitting.

### 3.3. Implementation

DeconV is implemented in Python with Pyro, a probabilistic programming language (Bingham et al. 2019), on top of PyTorch (Paszke et al. 2019), a deep learning framework. Probabilistic models defined by DeconV are fitted as described above using the stochastic gradient descent (SGD) based algorithm, Adam (Kingma and Ba 2014). In particular, Pyro’s clipped Adam optimizer is used with learning rate decay. This optimizer performs gradient clipping to prevent exploding gradients. With graphical processing unit (GPU)-acceleration enabled, deconvolution pipeline of DeconV can be sped up to 10-times. DeconV uses single-cell analysis framework, ScanPy (Wolf et al. 2018), to store, preprocess, and visualise both single-cell reference and bulk datasets.

### 3.4. Description of benchmarking datasets

DeconV was benchmarked on two datasets: Xin-dataset and PBMC-dataset. Xin-dataset is an artificial dataset popularized by the authors of MuSiC and utilized as a deconvolution benchmark in several publications (Wang et al. 2019; Menden et al. 2020, Dong et al. 2021). It consists of a single-cell reference from Segerstolpe et al. (Segerstolpe et al. 2016) and 18 samples of artificial pseudo-bulk generated from single-cell dataset (‘GSE81608’ Xin et al. 2016) containing 12 samples from healthy individuals and 6 samples from diseased individuals (type 2 diabetes). Since the bulk samples were composed of four cell types, namely alpha, beta, delta, and gamma, other cell types were filtered out from the single-cell reference, resulting in about 1000 cells remaining in the reference. Although the authors of MuSiC hypothesised that using multi-subject reference, i.e. sample- or condition-specific cell type reference, can improve the deconvolution accuracy, here we show that a high accuracy in deconvolution can be achieved without explicitly separating the cell type reference by condition.

The second dataset consists of a single-cell reference from a well known 3k PBMC-dataset by 10x Genomics. The bulk counterpart includes 18 samples of real bulk gene expression obtained from NCBI GEO archive ‘GSE107011’ (Monaco et al. 2019) where the ground truth cell type proportions were experimentally validated with flow cytometry. In contrast to the benchmark presented in the Scaden publication (Menden et al. 2020), where the same dataset was used as a benchmark with several single-cell references, we opted to use only a single reference with some modifications in cell type annotation. We also included the dendritic cells (DCs) into the annotation as those were present in the bulk samples, although in small quantity (0-2%). In addition, we removed ‘unknown’-annotation as this was consisting mostly of DCs prior to the changes. The annotation is based on several marker genes (**Table S2**) based on Seurat’s (Satija et al. 2015) guided clustering tutorial: https://satijalab.org/seurat/articles/pbmc3k_tutorial. We believe that using only a single reference for deconvolution better reflects real-world scenarios, due to the scarcity of relevant and high-quality single-cell references. In addition, integration of multiple single-cell datasets is often challenging due to many technical factors of sequencing, such as experimental design, sequencing technology and sequencing depth.

### 3.5. Execution of benchmarks

Deconvolution of two datasets described in previous section were carried out with DeconV and three methods from literature, namely CIBERSORTx (Newman et al. 2019), MuSiC (Wang et al. 2019), and Scaden (Menden et al. 2020). Scaden and MuSiC were executed with scripts based on examples provided by the developers, while CIBERSORTx was accessed through a web-application at https://cibersortx.stanford.edu. Default parameters were used for all methods, except for Scaden where the number of training samples was increased to 32,000 from 1,000, consistent with authors benchmarks on similar datasets. Reproducible deconvolution pipeline for benchmarks, including preprocessing, annotation, deconvolution, and plotting, are available from https://github.com/lutrarutra/deconv/tree/benchmark.

## 4. Discussion

DeconV is using a probabilistic approach that is an efficient way to solve the deconvolution problem . We assessed the accuracy on two datasets and compared the results with three methods from the literature. We showed that DeconV achieves state-of-the-art accuracy in deconvolution while improving the interpretability of the model and the results. This is achieved both by being able to extract the learned parameters of the model for each distribution and by giving confidence interval with the estimation of cell-type proportions.

The large discrepancies between real and *in silico* bulk dataset benchmark (**Figure S1**) pinpoints the urge to stop using artificial dataset for developing and benchmarking deconvolution methods. MuSic is a good study case as it performs well *in silico* but poorly with real data (**Table 1**). Further exploration on the fundamental differences between real and pseudo-Bulk RNA-Seq are needed to generate better *in silico* datasets of bulk RNA-Seq simulated from scRNA-seq. This would ease the development and benchmark of future deconvolution methods. An alternative is the generation of additional datasets as in Monaco et al. 2019, profiling the same samples with scRNA-seq and bulk RNA-seq. This approach would provide valuable resources for the field of deconvolution, which is currently experiencing a significant deficit preventing the production of large-scale benchmarks. Within DeconV, the systematic superiority of zero-inflated distribution over their regular counterparts show the usefulness of modelling with zero-inflation despite the recent controversies about this topic (Jiang et al. 2022). Furthermore, it is noteworthy that the ZINB distribution demonstrates superior performance with the PBMC dataset. This observation provides additional evidence supporting the efficacy of ZINB as a viable model for scRNA-seq data analysis.

The results from the PBMC dataset also highlight the problems of using coefficient of correlation as a measure of deconvolution accuracy. The results of gamma-model (**Figure 3c**) were reasonable: all cell type proportions were correctly estimated to fall on the lower range of the scale (*<* 40%). Whereas, a low coefficient of correlation (0.05) implies poor results. In addition, high correlation did not necessarily translate to a smaller error and vice versa. For example, Scaden exhibited better correlation but also a higher error than DeconV (**Table 1**). We recommend using additional error measures, such as the ones proposed in this work (ie. RMSE), to assess correlation in future benchmarks.

The source code and example scripts for DeconV are available on Github at github.com/lutrarutra/deconv. Reproducible benchmarks are available from github.com/lutrarutra/deconv/tree/benchmark.

## 5. Supplementary

### 5.1. Supplementary method

The justification that bulk RNA-Seq is still more common than scRNA-Seq is justified by the number of results for the following queries on PubMed:

- 84,870 results for the query *(“RNA-Seq” OR “RNA sequencing” OR “transcriptome sequencing”) NOT (“single-cell RNA sequencing” OR “scRNA-Seq” OR “single-cell transcriptomics”)*
- 14,225 results for the query *“single-cell RNA sequencing” OR “scRNA-Seq” OR “single-cell transcriptomics”*

## 5.2. Supplementary figures

**Table S1:** Five configurations for modeling dropouts. In shared dropout models, dropout coefficient is same for all cell types while in separate dropout models dropout can vary between cell types.

| Name | Dropout type in scRNA-seq | Dropout in bulk RNA-seq |
| --- | --- | --- |
| no dropout | None | False |
| shared | shared | True |
| separate | separate | True |
| shared, no bulk | shared | False |
| separate, no bulk | separate | False |

**Table S2:** Marker genes used for annotation of single-cell reference.

| Cell Type | Markers | Assigned Label |
| --- | --- | --- |
| Naive CD4+T | IL7R, CCR7 | T CD4 |
| Memory CD4+T | IL7R, S100A4 | T CD4 |
| CD14+ Mono | CD14, LYZ | Monocytes |
| FCGR3A+ Mono | FCGR3A, MS4A7 | Monocytes |
| B-cells | MS4A1 | B-cells |
| T CD8 | CD8A | T CD8 |
| NK | GNLY, NKG7 | NK |
| Dendritic Cells | FCER1A, CST3 | DCs |

**Figure S1:**
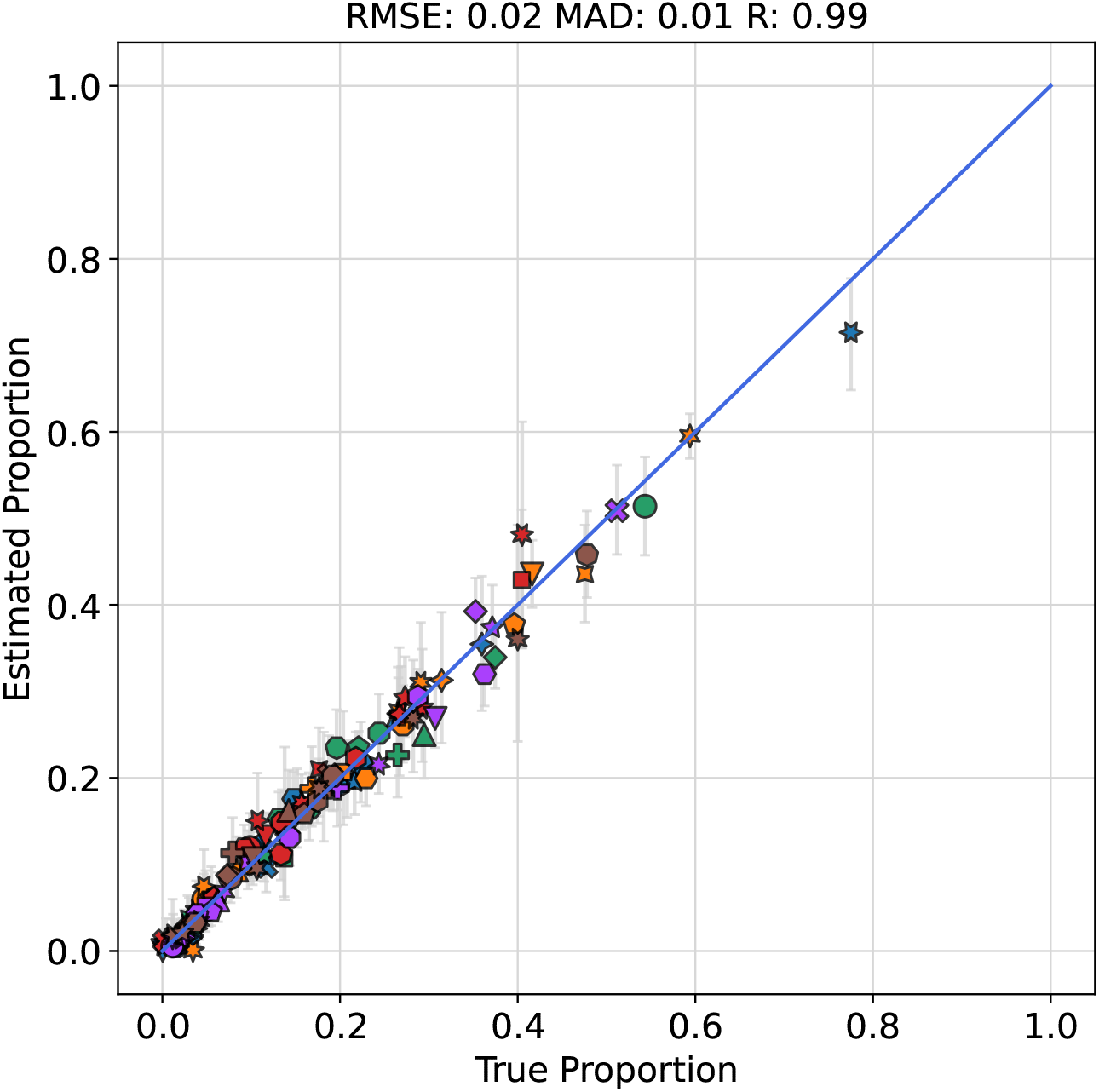
Deconvolution of a synthetic PBMC-dataset consisting of 20 pseudo-bulk samples. Pseudo-bulk samples were generated by drawing cell type proportions from dirichlet distribution and summing gene-expression of 1000 cells. Each point corresponds to a proportion of a sample. Color denotes cell types while shape denotes samples. For deconvolution, negative-binomial model was used.

**Figure S2:**
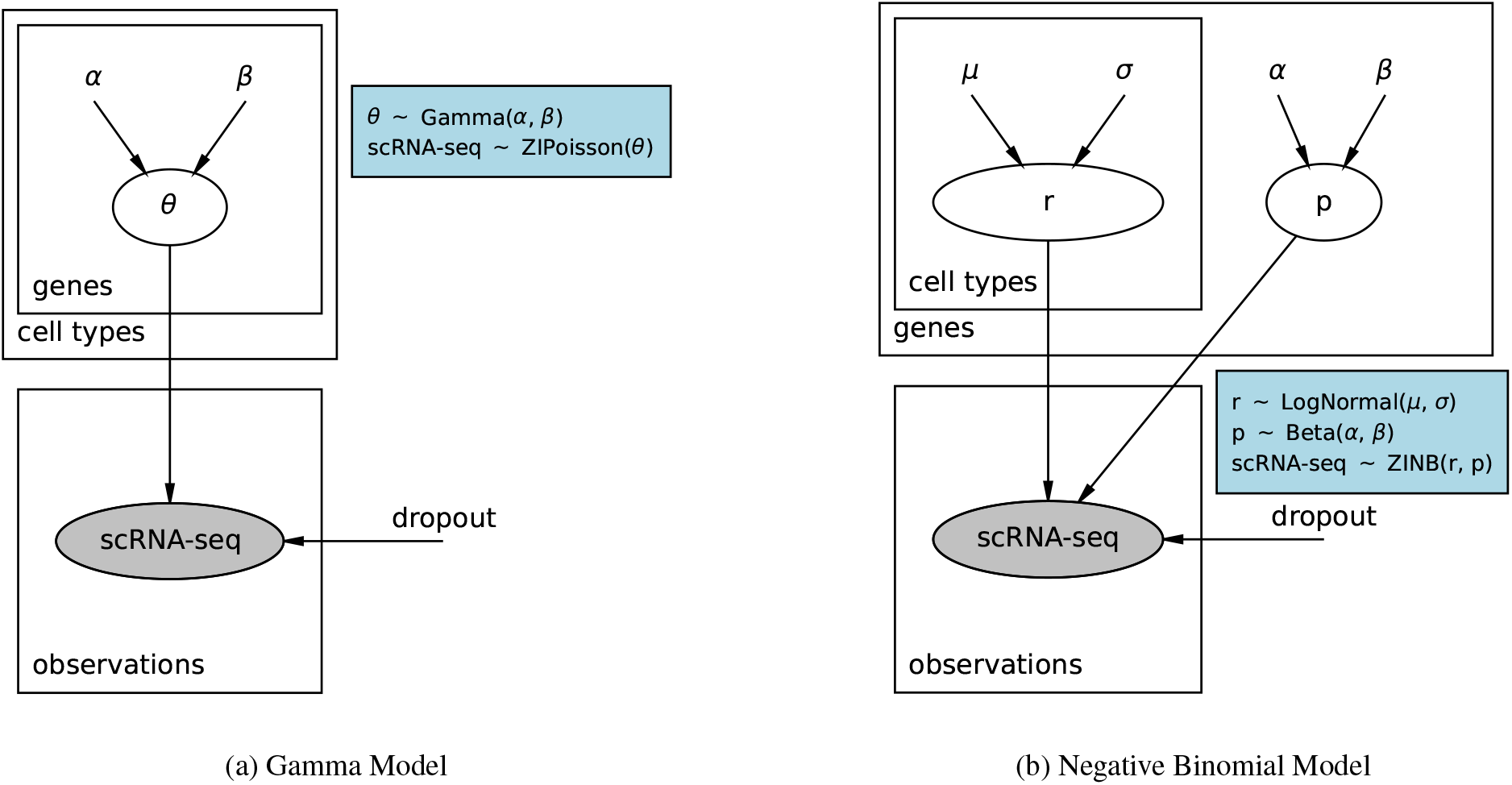
Graphical illustrations of the probabilistic cell type reference models defined by DeconV. Variables inside circles are sampled from distributions, highlighted in blue boxes. Grey circles indicate observed data while circles with white background are latent parameters of the models. Variables outside any circles are learnable parameters that are fitted so that the likelihood of the observed data is maximized.

